# gplasCC: classification and reconstruction of plasmids from short-read sequencing data for any bacterial species

**DOI:** 10.1101/2024.11.28.625923

**Authors:** Julian A. Paganini, Jesse J. Kerkvliet, Gijs Teunis, Oscar Jordan, Nienke L. Plantinga, Rodrigo Meneses, Rob J.L. Willems, Sergio Arredondo-Alonso, Anita C. Schürch

## Abstract

Plasmids play a pivotal role in the spread of antibiotic resistance genes. Accurately reconstructing plasmids often requires long-read sequencing, but bacterial genomic data in publicly accessible repositories has historically been derived from short-read sequencing technology. We recently presented an approach for reconstructing *Escherichia coli* antimicrobial resistance plasmids using Illumina short reads. This method consisted of combining a robust binary classification tool named plasmidEC with gplas2, which is a tool that makes use of features of the assembly graph to bin predicted plasmid contigs into individual plasmids. Here, we developed gplasCC, a plasmidEC-simplification, capable of classifying plasmid contigs using Centrifuge databases. We have developed seven plasmidCC databases in addition to the database for *E. coli*: six species-specific models (*Acinetobacter baumannii*, *Enterococcus faecium*, *Enterococcus faecalis*, *Klebsiella pneumoniae*, *Staphylococcus aureus* and *Salmonella enterica*) and one species-independent model for less frequently studied bacterial species. We combined these models with gplas2 (now, gplasCC) to reconstruct plasmids from more than 100 bacterial species. This approach allows comprehensive analysis of the wealth of bacterial short-read sequencing data available in public repositories and advance our understanding of microbial plasmids.

## INTRODUCTION

Antimicrobial resistance (AMR) is a major threat to human health worldwide. Estimates indicate 1.27 million deaths were attributable to bacterial AMR in 2019 alone^1^. Moreover, the number of infections caused by resistant bacteria increases yearly. In recent years, only a few new antibiotics have been approved by the FDA^2^, and their use is recommended for a limited number of clinical scenarios^3^. Although alternative approaches to treating bacterial infections are being explored, their effects have seen limited clinical application to date^4–7^. Consequently, it will take several years for these methods to become commonplace to treat bacterial infections. In this perspective, limiting the dissemination of resistance among bacteria is the key to preventing a further AMR crisis.

The spread of AMR is a complex phenomenon that depends on various factors. However, it has become increasingly clear that plasmids play a central role in this process. Plasmids are mobile genetic elements (MGE) that frequently carry AMR genes and can be transferred between bacteria by diverse mechanisms^8–12^. Additionally, plasmids play an important role in outbreaks in clinical settings involving multiple bacterial species^13–17^. Therefore, accurate, high-throughput plasmid identification and tracking is becoming increasingly important and knowledge on plasmid diversity and evolution is urgently needed.

Next-generation sequencing (NGS) platforms offer powerful tools for large-scale bacterial genome research. Despite the technological advancements of long-read technologies, which enable the sequencing of complete bacterial genomes^18,19^, Illumina short-read remains the most widespread sequencing method, both historically and currently. As of December 2023, the Sequence Read Archive (SRA) contained more than 2.3 million DNA sequences belonging to bacterial isolates and 97.8% of these were obtained using short-reads technology (Figure 1). However, repeated elements commonly present on plasmids complicate plasmid assembly by short-read data alone^20^. Therefore, specialized post-assembly methods for reconstructing plasmids from short-read data are needed.

**Figure 1.**
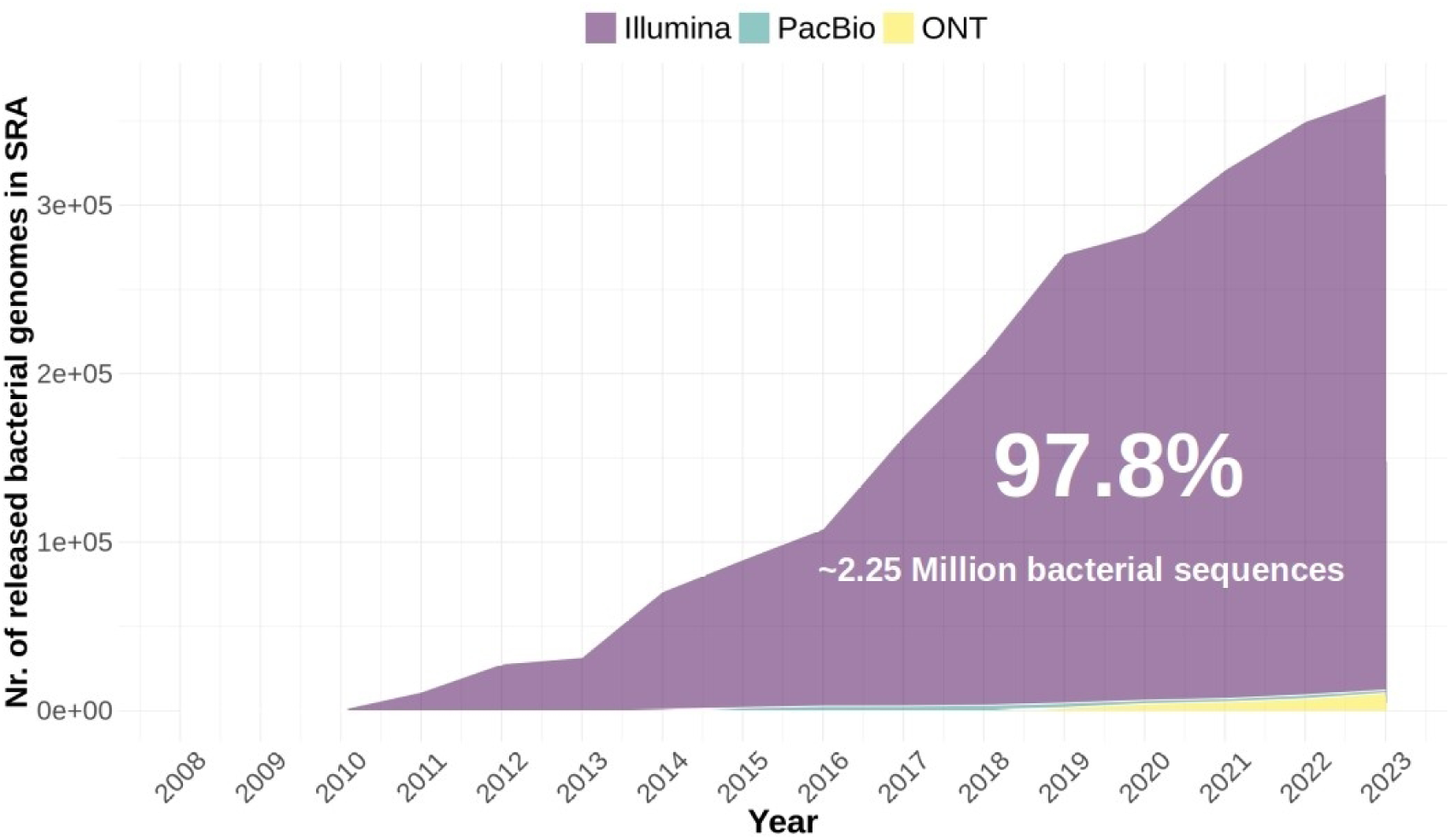
Number of genomes sequenced by different NGS technologies, publicly available in the Sequence Read Archive (SRA) by 31st December 2023. 97.8% of genomes were sequenced with Illumina short-read technology. released genomes per species

Recently, we have developed a method for reconstructing individual *Escherichia coli* plasmids from short reads ^21^. In this method, nodes in the assembly graph were initially classified as plasmid- or chromosome-derived using plasmidEC, an ensemble classifier that combines the output of three existing binary classification tools^22–24^. After this initial classification step, we used gplas2^21^ to bin plasmid nodes into individual plasmid bins based on sequence coverage similarities and assembly-graph connectivity. Our method outperformed the commonly used tool MOB-suite^25^, especially when reconstructing plasmids carrying antibiotic-resistance genes (ARGs).

In this study, we aimed to improve the classification and reconstruction of plasmids in multiple additional bacterial species using short-read data. To improve plasmid-fragment classification, we developed plasmidCC (plasmid Centrifuge Classifier), an adapted, simplified version of plasmidEC. This tool uses Centrifuge databases^26^ for plasmid classification. We constructed species-specific Centrifuge databases for seven common human pathogens (*Acinetobacter baumannii*, *Enterococcus faecium*, *Enterococcus faecalis*, *Escherichia coli, Klebsiella pneumoniae*, *Salmonella enterica*, and *Staphylococcus aureus*)^1,27,28^ that are sequenced frequently and are consequently well-represented in public databases^29^. In addition, we developed a species-independent Centrifuge classifier to predict plasmids from 106 different less frequently represented species in sequencing databases.

Furthermore, to improve plasmid reconstruction, we developed a new, lightweight version of gplas2 called gplasCC. This improved version of gplas has direct integration with plasmidCC to combine the plasmid classification and reconstruction steps into one step. Furthermore, the improved gplas algorithm assigns repeated sequences to their appropriate plasmid bins, whereas gplas2 would omit assigning these sequences to a plasmid bin. We applied gplasCC to the output of plasmidCC to reconstruct individual plasmids from the species mentioned above. To gauge the effectiveness of our approach, we compared its performance to the results of the popular plasmid reconstruction tools MOB-suite and plasmidSPAdes^25,30^.

## MATERIAL AND METHODS

### Querying the SRA database

To create a general overview of the contents of the SRA database per sequencing technology, we retrieved the metadata associated with bacterial whole genome sequences. We used the esearch function of the package Entrez Direct (v13.3)^31^ to query the database and the efetch function to extract the metadata. The search terms included in this search are described in the Supplementary methods. The following search terms were included:

esearch -db sra -q ‘(“Bacteria”[Organism] OR “Bacteria Latreille et al. 1825”[Organism]) AND “platform illumina”[Properties] AND (cluster_public[prop] AND “biomol dna”[Properties] AND “strategy wgs”[Properties])’ | efetch -format summary

esearch -db sra -q ‘(“Bacteria”[Organism] OR “Bacteria Latreille et al. 1825”[Organism]) AND “platform illumina”[Properties] AND (cluster_public[prop] AND “biomol dna”[Properties] AND “strategy wgs”[Properties])’ | efetch -format runinfo

The platform term was varied accordingly to include the three sequencing technologies ‘illumina’, ‘Oxford Nanopore’ and ‘PacBio SMRT’.

### Development of plasmidCC models

PlasmidCC classifies plasmid sequences using custom-built Centrifuge databases. Centrifuge^26^ is a taxonomy classifier for metagenomics reads. We adapted this tool to function as a binary classifier of short-read whole genome sequencing contigs by building databases containing complete bacterial sequences labelled as either plasmid or chromosome, similar to what has been described for PlaScope^24^. In contrast to PlaScope, plasmidCC calculates the per-contig proportion of plasmid-based and chromosome-based hits and uses a threshold of 70% plasmid-based hits for plasmid classification and a threshold of 30% of chromosome-based hits for chromosome classification. Contigs with classification proportions between 30% and 70% are labelled ‘unclassified’.

We have built seven species-specific databases for classification of contigs of *K. pneumoniae*, *A. baumannii*, *S. aureus*, *S. enterica*, *E. faecium* and *E. faecalis*. Additionally, we have implemented the *E. coli* database developed by PlasCope^24^, and a species-independent database, hereafter called the General database, using all available genomes from any bacterial species, aside from the seven aforementioned species. Genomes were included if they were published to RefSeq^32^ before 2022.

### Description of gplasCC

gplasCC is an extension of the previously published gplas algorithm^21,33^ and combines the plasmid classification of plasmidCC with the plasmid reconstruction of gplas. gplasCC uses plasmidCC (version 1.0.0) to classify the nodes in a given short-read assembly graph as plasmid- or chromosome-derived. Next, to quantify the expected variance in coverage between plasmid- and chromosome-derived nodes, the distribution of coverage of the chromosome-predicted nodes is calculated. In turn, the standard deviation of this distribution is retained as a threshold to determine which nodes have a similar coverage. Next, random walks, starting from plasmid-classified nodes are performed, aiming to connect plasmid-classified nodes with a similar coverage using a greedy approach. After 20 iterations of this step, gplasCC constructs undirected, weighted network graphs connecting plasmid nodes based on co-occurrence of nodes in multiple random walks. These weighted network graphs are subjected to three community partitioning algorithms (Louvain, Leading-eigen and Walktrap)^34,35^. When the modularity of the created community-subnetworks exceeds the modularity threshold (0.2 by default) in at least two of the algorithms, the subnetwork is split according to the partitioning with the highest modularity.

If a subset of plasmid-classified nodes remains unbinned, gplasCC employs bold mode, repeating the random walks with relaxed coverage constraints.

In earlier iterations of gplas^21,33^, a node was defined as repeated if it had more than one incoming or outgoing edge. Such nodes were excluded from plasmid reconstructions. To address this limitation, gplasCC introduced a step that systematically assigns this set of repeated nodes to the most likely plasmid bin or bins.

In gplasCC, plasmid bins are constructed as a set of nodes that are not present in any other plasmid bin and are not included in the set of repeated nodes. Nodes that are neither in any plasmid bin nor in the set of repeated nodes are classified as chromosomal nodes. Repeat assignment is performed based on these plasmid bins and the collection of chromosomal nodes. Each repeat node is a seed for a set of short random walks (up to 5 nodes) through the graph in both directions. The walk terminates either when a non-repeated node is encountered (i.e. a node in a plasmid bin or a chromosomal node) or after five steps.

A walk in a particular direction is deemed valid if either 1) it concluded at a non-repeated node after one step, or 2) at least one walk in the opposite direction concluded at a node with the same bin membership.

For each repeat node, we count the endpoints of the valid walks and establish a ranking based on the relative frequency of the (plasmid or chromosomal) bins these endpoints are part of. The repeated node is assigned to the first bin in the ranking if it meets the following coverage requirement:

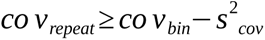

where *cov*_repeat_ is the sequencing coverage of the repeated element, *cov_bin_* represents the average coverage of all unitigs composing that bin and *s^2^_cov_* is the variance in sequencing coverage estimated in an earlier step of the gplas algorithm^33^.

Upon assignment, the repeat node is associated with this bin, and its coverage is updated by subtracting the group coverage of the newly assigned bin from the coverage of the repeated node. This process repeats for subsequent bins in the ranking until the coverage requirement is no longer met. Each repeat can only be assigned once to each bin.

### Benchmarking of plasmidCC and gplasCC

#### Compiling benchmarking datasets

PlasmidCC was benchmarked against RFPlasmid^23^, Platon^22^ and (for *E. faecalis* and *E. faecium*) mlplasmids^36^. The performance of gplasCC was compared to that of plasmidSPAdes and MOB-suite.

To validate the performance of these tools, we compiled a benchmarking dataset containing plasmids belonging to seven common human pathogens that are abundant in sequence databases as well as plasmids belonging to less commonly sequenced species. To exclude genomes used in the development of MOB-suite, we included only those genomes from the common human pathogens that were uploaded to RefSeq after May 2020. For less frequently researched species in our general dataset, we included genomes uploaded in 2022.

#### Genome assembly and quality trimming

Complete genomes and corresponding short reads were downloaded from RefSeq and SRA using ncbi-genome-download (v0.2.10) (https://github.com/kblin/ncbi-genome-download) and SRA tools (v2.10.9) (https://github.com/ncbi/sra-tools), respectively.

Illumina raw reads were trimmed using trim-galore (v0.6.6) (https://github.com/FelixKrueger/TrimGalore) to remove adapter contamination and bases with a phred quality score below 20. Unicycler^37^ (v0.4.8) was then applied to perform de novo assembly with default parameters.

### Exploring the plasmid diversity in benchmarking datasets

To establish an overview of the plasmid diversity in both the dataset used for the database construction, as well as the benchmarking dataset, we used Mash^38^ v2.2.2 (k = 21, s = 10,000) to estimate the pairwise k-mer distances of all complete plasmid sequences of *A. baumannii, K. pneumoniae, E. faecium, E. faecalis, S. aureus S. enterica*. We clustered the obtained distances using the t-distributed stochastic neighbour embedding algorithm (t-SNE)^39^ with a perplexity value of 30. Data points representing individual sequences of plasmids were coloured if they were part of the benchmarking dataset. A similar analysis for the *E. coli* dataset can be found in ^40^.

#### Sub-classifying plasmids according to their length

Plasmids were classified as large or small based on size distributions for each species. For *E. coli* ^40^, *E. faecium, E. faecalis, K. pneumoniae* and *S. enterica* cut-off values of 18,000 bp were selected; while for *A. baumannii* and *S. aureus* cut-off values were 30,000 bp and 8,000 bp, respectively.

#### Determining the origin of all assembled contigs

After assembly, resulting contigs were labelled as chromosome- or plasmid-derived by alignment to their corresponding complete genomes using QUAST (v5.0.2)^41^. Only contigs larger than 1,000 bp with an alignment of at least 90% of the contig length were considered. Of those, contigs aligning to both the chromosome and plasmidome (ambiguously aligned contigs) were considered repeated and not used when calculating reconstruction metrics. Contigs larger than 1kb that aligned to multiple locations in the genomes were labelled as repeats. These contigs were classified as ‘ambiguous’ by QUAST.

### Evaluating binary classification

We evaluated the performance of plasmidCC by comparing each contig’s prediction to its actual class. All ‘unclassified’ predictions were considered chromosomal. The predictions were categorised as follows: True Positives (TP, correctly predicted plasmid), True Negatives (TN, correctly predicted chromosome), False Positives (FP, incorrectly predicted plasmid) and False Negatives (FN, incorrectly predicted chromosome). We evaluated the global performance of the tools with the following metrics:

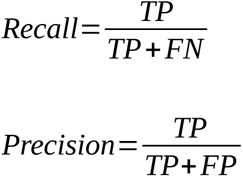

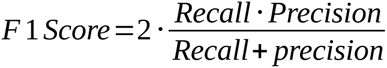

### Evaluating plasmid reconstruction tools

To reconstruct individual plasmids, gplasCC was run using the output of plasmidCC on all benchmarking plasmids.

To evaluate bins created by MOB-suite, plasmidSPAdes and gplasCC, we used QUAST (v5.0.2) to align the contigs of each bin to the corresponding complete reference genome. To evaluate the performance of the plasmid reconstruction tools, we calculated metrics often used in metagenomics to compare genome binning^42,43^. We calculate a per-plasmid completeness score and a per-bin accuracy score, as defined below, to assess the tool’s ability to create bins containing complete plasmids without contamination of other plasmid or chromosomal contigs. We also evaluated the number of reference plasmids detected by each tool. We considered a reference plasmid detected when at least a single contig of the plasmid was included in the predictions.

Additionally, to assess the ability of the reconstruction tools to create the correct number of bins equal to the number of plasmids, we calculated the binning rigour. All calculated metrics are specified below.

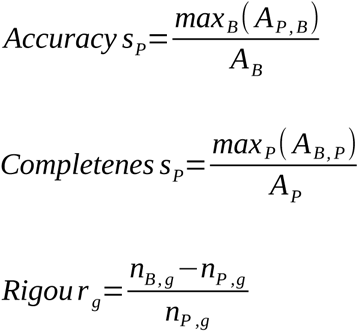

Where *A* is the alignment length in base pairs (bp) for each plasmid *P* and bin *B*, and *A_B_* and *A_P_* are the total length in bp for each bin and plasmid, respectively. Binning rigour for genome *g* was calculated by subtracting the number of plasmids *n_P_* for genome g from the number of bins *n_B_* produced for genome *g*, divided by the number of plasmids in the genome.

We reported the chromosome contamination per bin to evaluate the incorrect inclusion of chromosome-derived contigs in the bins as the fraction of bp in a bin that aligned to a chromosome.

To systematically assess the performance of each tool in generating high-quality bins we categorized each bin into one of five categories based on their accuracy and the completeness of the largest plasmid fraction in the bin. Bins classified as ‘very high’ quality contained a plasmid fragment with a minimum completeness of 0.90 and a contamination level of no more than 10% (accuracy ≥ 0.9). Those designated as ‘high’ quality had a completeness exceeding 0.7 and a contamination level of no more than 30% (accuracy ≥ 0.7). ‘Moderate’ quality bins were characterized by a completeness greater than 0.5 and an accuracy greater than 0.5. We categorized bins with a completeness exceeding 0.3 and an accuracy of at least 0.3 as ‘low’ quality, in contrast to those with less than 0.3 completeness and accuracy below 0.3, which were deemed ‘very low’ quality.

#### ARG Predictions

ARGs in completed genomes and plasmid bins were predicted by running Abricate (v1.0.1) (https://github.com/tseemann/abricate) with the Resfinder^44^ database (indexed on 19 April 2020) using reference plasmids as query, with 80% identity and coverage cut-off.

#### Evaluating the capacity of the tools to correctly assign ARGs to plasmid predictions

All ARGs assigned to plasmid bins were classified as plasmid-borne (Detected) or chromosomal (Contamination) by determining the origin of the contig carrying the ARGs, as detailed above. We obtained the number of undetected ARGs by subtracting the number of detected ARGs from the total number of predicted plasmid-borne ARGs in reference genomes.

#### Evaluation of the gplas2 repeat resolution algorithm

To evaluate the performance of the repeat resolution algorithm of gplasCC, we compared each repeat in a given isolate to the true class of that repeat. We considered an assignment a true positive (TP) when the repeat aligned to at least one plasmid and was assigned to at least one plasmid bin. We considered an assignment a true negative (TN) when the repeat aligned to one or more chromosomal regions and was not assigned to a plasmid bin. We considered a repeat a false positive (FP) when the repeat aligned exclusively to one or more chromosomal regions, but it was assigned to a plasmid bin, and we considered a repeat a false negative (FN) when it aligned to at least one plasmid but was not assigned to a plasmid bin.

## RESULTS

### Construction of Centrifuge databases

Eight custom Centrifuge databases were constructed using complete genome assemblies in the RefSeq database. For the seven species-specific databases, a total of 3,687 genomes were used, whereas the general database was constructed using 25,735 genomes. Distribution of genomes per database can be found in Figure 2A. The distribution of species in the general database can be found in Figure 2B.

**Figure 2.**
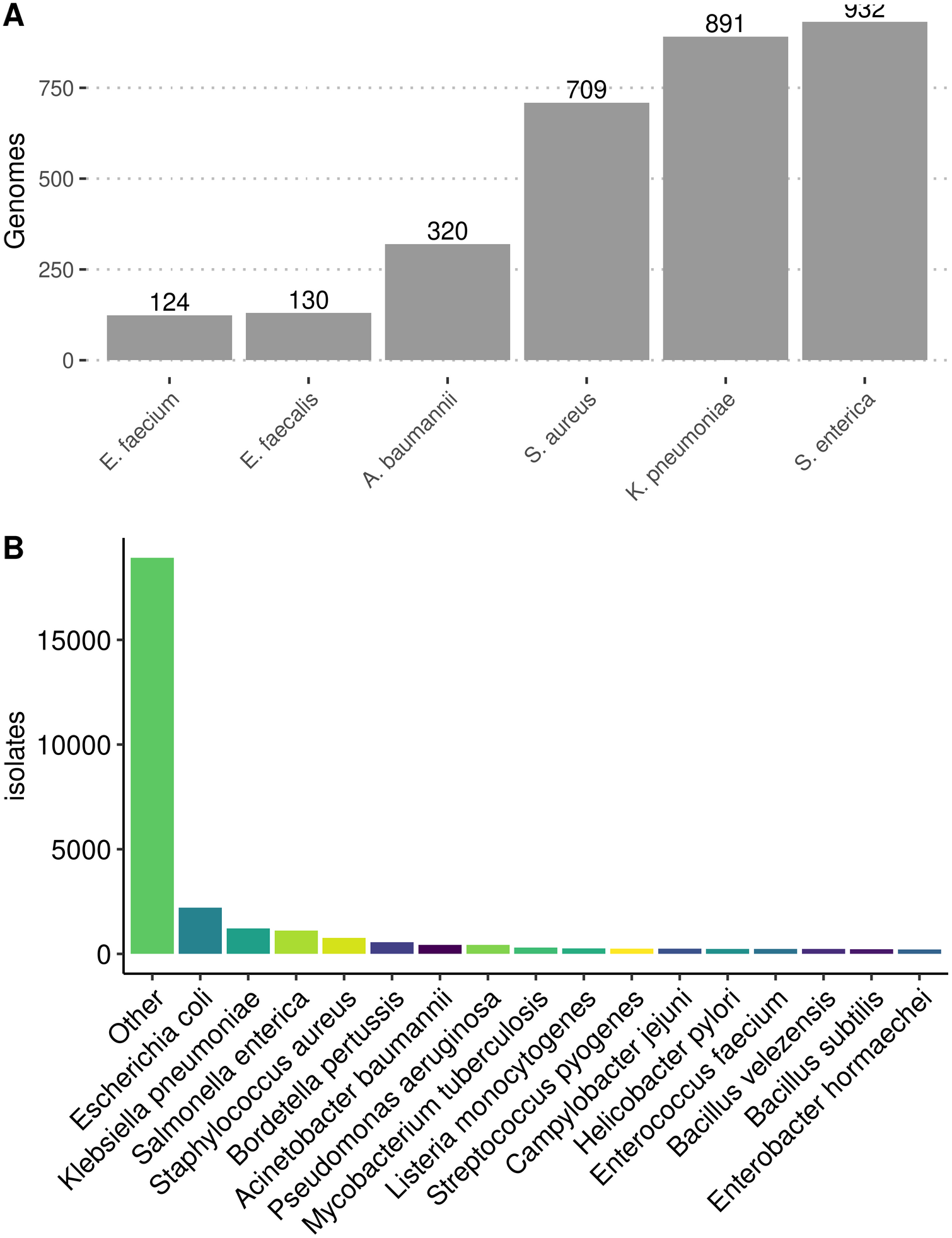
**A)** Number of genomes included in the construction of species-specific Centrifuge databases. **B)** Genomes included in the construction of the species-independent Centrifuge database. The ‘Other’ category includes genomes from species that are contain less than 200 genomes in RefSeq.

### Evaluating binary classification

To compare the performance of plasmidCC against multiple existing classification tools we used a benchmarking dataset consisting of 981 genomes, harbouring a total of 2620 plasmids. A complete list of genomes used in the benchmarking dataset can be found in Supplementary Data 1. Of these, 710 genomes and 2032 plasmids belonged to species highly abundant in the public databases with high clinical relevance (*A. baumannii, E. coli, E. faecalis, E. faecium, K. pneumoniae, S. aureus, S. enterica*). The remaining 277 genomes and 609 plasmids belonged to 107 species, less frequently represented in databases (Supplementary Figure S1). Plasmids from *A. baumannii, E. faecalis, E. faecium, K. pneumoniae, S. enterica* and *S. aureus* captured most of the available plasmidome diversity for each species (Supplementary Figure S2).

PlasmidCC achieved the highest F1-Score values when classifying contigs from six species and the species-agnostic dataset (Figure 3, Table 1). In the case of *S. enterica*, plasmidCC was outperformed by a majority voting of Platon, plasmidCC and RFPlasmid. None of the individual tools outperformed plasmidCC. PlasmidCC had the highest precision in all species except *K. pneumoniae* and the highest recall for three species-specific datasets (*A. baumannii, E. coli, and K. pneumoniae*) and when using the general database, although the difference was minimal for *E. faecium* (difference = 0.0017).

**Figure 3.**
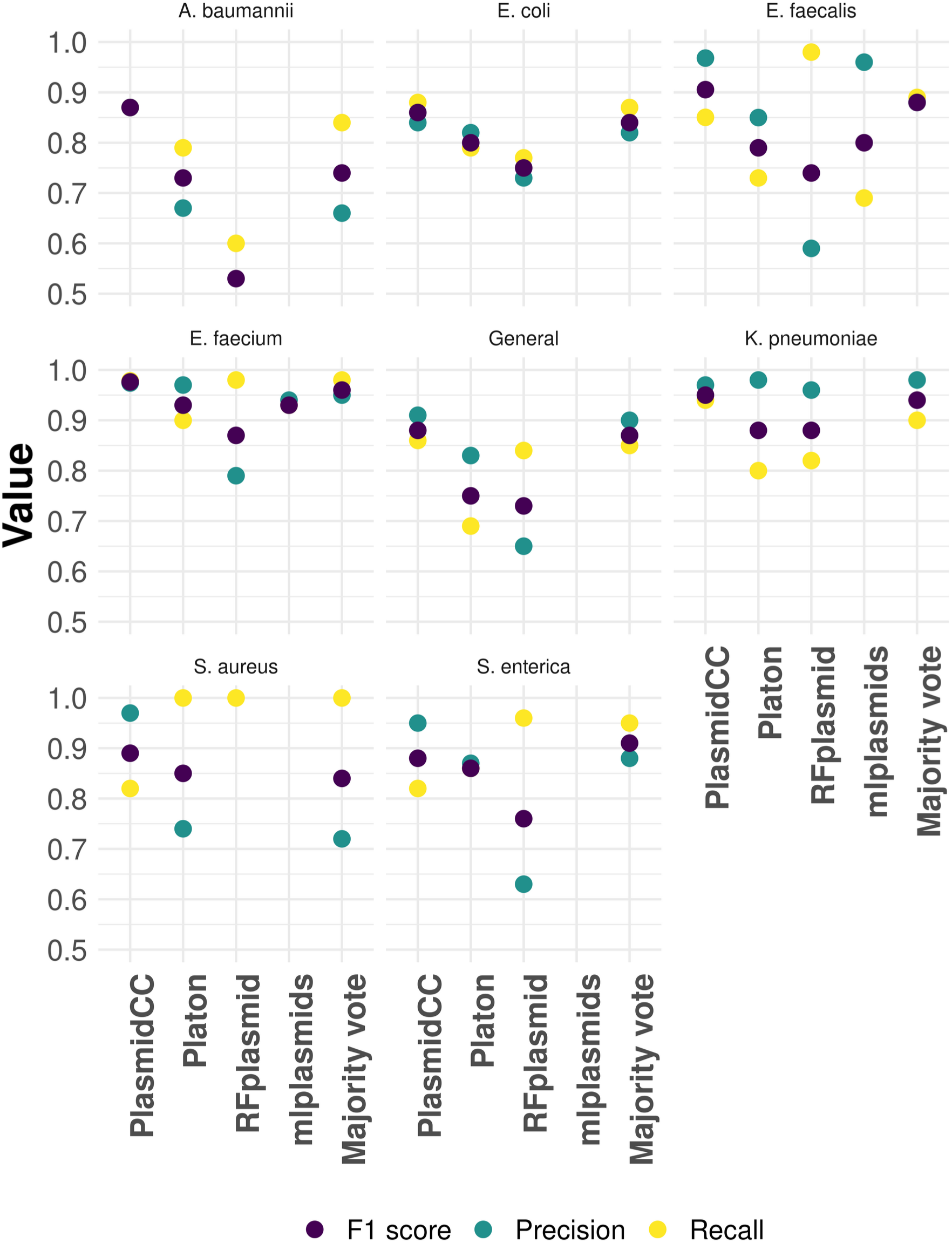
Performance of different individual binary classification tools and the results of a majority voting classifier. The ‘General’ category corresponds to species less frequently represented in databases (See Supplementary Figure S1B for a summary of included species). Mlplasmids was only used on species with a suitable model (*E. faecium* and *E. faecalis*).

**Table 1.** Classification metrics recall, precision and F1-score per species and per tool. Bold values indicate highest value.

| Species | Tool | F1 score | Precision | Recall |
| --- | --- | --- | --- | --- |
| *A. baumannii* | **plasmidCC** | **0.87** | **0.87** | **0.87** |
|  | RFplasmid | 0.53 | 0.47 | 0.6 |
|  | Platon | 0.73 | 0.67 | 0.79 |
|  | Majority voting | 0.74 | 0.66 | 0.84 |
| *E. coli* | **plasmidCC** | **0.86** | **0.84** | **0.88** |
|  | RFplasmid | 0.75 | 0.73 | 0.77 |
|  | Platon | 0.8 | 0.82 | 0.79 |
|  | Majority voting | 0.84 | 0.82 | 0.87 |
| *E. faecalis* | **plasmidCC** | **0.91** | **0.97** | **0.85** |
|  | RFplasmid | 0.74 | 0.59 | 0.98 |
|  | Platon | 0.79 | 0.85 | 0.73 |
|  | Majority voting | 0.88 | 0.88 | 0.89 |
|  | mlplasmids | 0.8 | 0.96 | 0.69 |
| *E. faecium* | **plasmidCC** | **0.98** | **0.97** | **0.98** |
|  | RFplasmid | 0.87 | 0.79 | 0.98 |
|  | Platon | 0.93 | 0.97 | 0.9 |
|  | Majority voting | 0.96 | 0.95 | 0.98 |
|  | mlplasmids | 0.93 | 0.94 | 0.93 |
| *K. pneumoniae* | **plasmidCC** | **0.95** | **0.97** | **0.94** |
|  | RFplasmid | 0.88 | 0.96 | 0.82 |
|  | Platon | 0.88 | 0.98 | 0.8 |
|  | Majority voting | 0.94 | 0.98 | 0.9 |
| *S. aureus* | **plasmidCC** | **0.89** | **0.97** | **0.82** |
|  | RFplasmid | 0.37 | 0.23 | 1 |
|  | Platon | 0.85 | 0.74 | 1 |
|  | Majority voting | 0.84 | 0.72 | 1 |
| *S. enterica* | plasmidCC | 0.88 | 0.95 | 0.82 |
|  | RFplasmid | 0.76 | 0.63 | 0.96 |
|  | Platon | 0.86 | 0.87 | 0.86 |
|  | **Majority voting** | **0.91** | **0.88** | **0.95** |
| *General* | **plasmidCC** | **0.88** | **0.91** | **0.86** |
|  | RFplasmid | 0.73 | 0.65 | 0.84 |
|  | Platon | 0.75 | 0.83 | 0.69 |
|  | Majority voting | 0.87 | 0.9 | 0.85 |

### Evaluation of the performance of gplasCC

We compared the performance of gplasCC against MOB-suite and plasmidSPAdes by applying all tools to the genomes of the benchmark dataset. Plasmid bins produced by each tool were aligned to the corresponding complete reference genomes and assessed using the metrics accuracy, completeness, binning rigour and relative bin quality. Furthermore, we evaluated the capacity of gplasCC to assign repeated sequences to their correct respective bins.

We split plasmids by size into ‘large’ and ‘small’ categories based on cut-offs obtained from the distribution of plasmid sizes shown in Supplementary Figure S3. In addition, we split large plasmids into large ARG plasmids and large non-ARG plasmids (Supplementary Table S1).

When reconstructing large ARG plasmids (n=650), gplasCC had a global median accuracy of 0.933 (IQR = 0.767-1.00), outperforming plasmidSPAdes (median=0.891,IQR=0.516 - 0.994), but falling second to MOB-suite (median=0.952, IQR=0.816-1.000) (Figure 4A and B). PlasmidSPAdes showed the highest completeness value (median=0.838, IQR=0.528-0.948), but had a high level of variation in completeness values, as indicated by the wide IQR. GplasCC had the second highest completeness value (median = 0.826, IQR = 0.656-0.928), followed by MOB-suite (median=0.778, IQR=0.516-0.923). Similar results were observed when reconstructing large plasmids without resistance genes (n=891) (Supplementary Figure S3).

**Figure 4.**
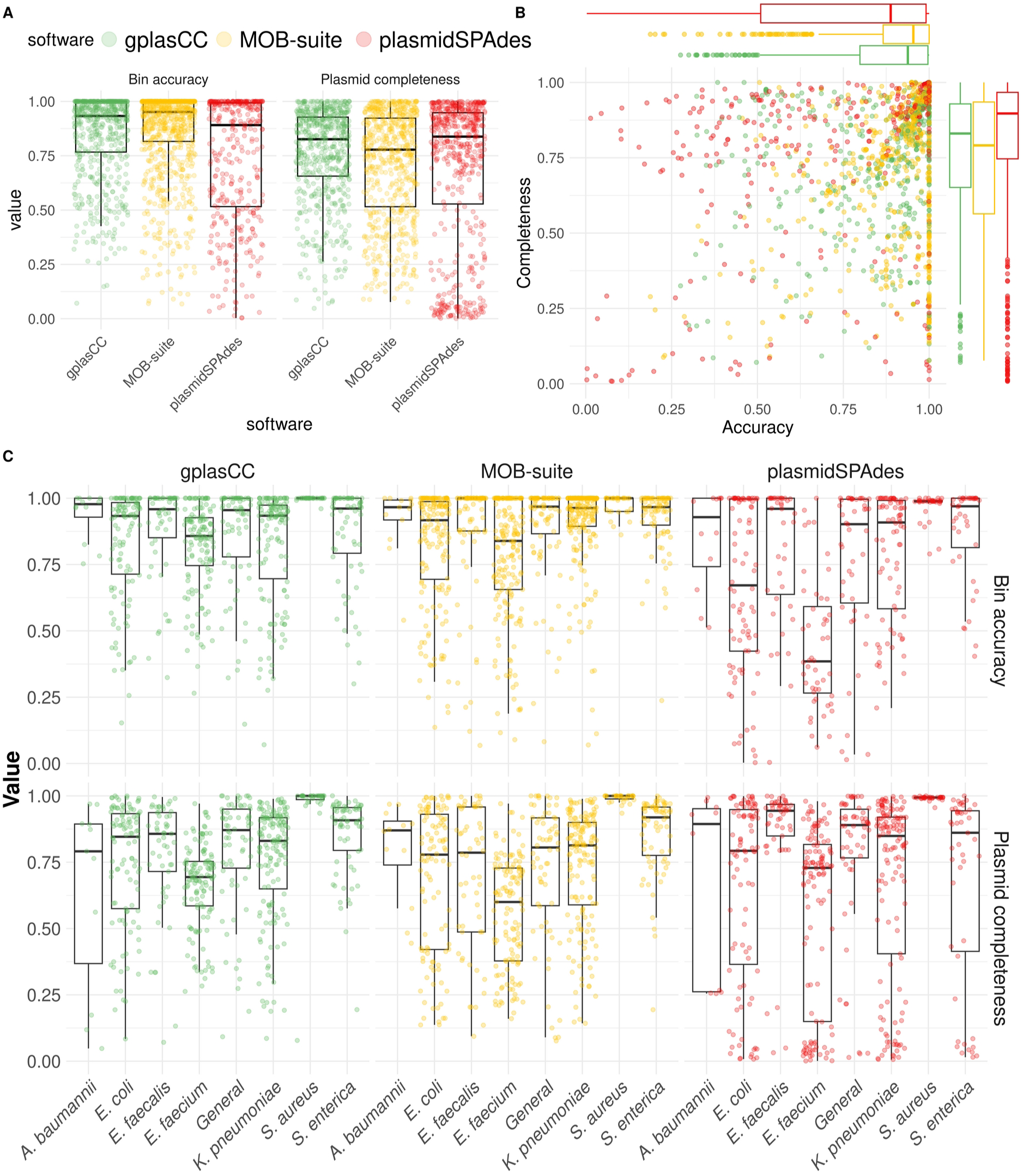
Reconstruction metrics of gplasCC, MOB-suite and plasmidSPAdes. **A)** Bin accuracy and plasmid completeness per tool, **B)** Plasmid completeness (y-axis) versus bin accuracy (x-axis) for large ARG plasmids from all species. **C)** Reconstruction metrics for detected large ARG plasmids per species (bottom).

Next, we analyzed the reconstruction of large ARG-plasmids for each species separately (Figure 4C, Supplementary Table S2). Compared to the other tools, gplasCC achieved the highest median completeness score for *E. coli* (0.846), and *S. aureus,* where two tools (gplasCC and MOB-suite) achieved a median completeness value of 1.000. GplasCC achieved the second highest median completeness values for four species-specific datasets (*E. faecalis, E. faecium, K. pneumoniae, S. enterica*) and the general dataset. For all but one of these cases gplasCC fell second to plasmidSPAdes. The exception was *S. enterica* where both plasmidSPAdes and gplasCC were outperformed by MOB-suite. *A. baumannii* is the only species where gplasCC scored the lowest median completeness, followed by MOB-Suite and plasmidSPAdes.

Furthermore, gplasCC had the highest median accuracy values for four species-specific datasets (*A. baumannii, E. coli, E. faecium, S. aureus*). PlasmidSPAdes scored the lowest median accuracy for all species, except for *E. faecalis* where gplasCC scored the worst and MOB-suite performed best, although the difference between gplasCC and plasmidSPAdes was minimal (0.959 and 0.960).

We further measured the tools’ ability to create the correct number of bins per plasmid by calculating per-isolate binning rigour for each tool (Figure 5A and B). Rigour values above zero indicate that a tool produced too many bins relative to the number of plasmids in a genome. In contrast, values below zero indicate too few bins relative to the number of plasmids. Calculating the rigour for each genome showed that, globally, plasmidSPAdes had a higher abundance of below zero rigour values. On the other hand, MOB-suite showed a higher abundance of above-zero rigour, as indicated by the peaks in distribution (Figure 5B). We observed the most notable shifts in the distribution of rigour in *E. faecalis, E. faecium and K. pneumoniae* for plasmidSPAdes and MOB-suite. A notably different distribution of rigour was observed for gplasCC in *E. faecalis* and *E. faecium*, where gplasCC tended to have lower rigour values.

**Figure 5.**
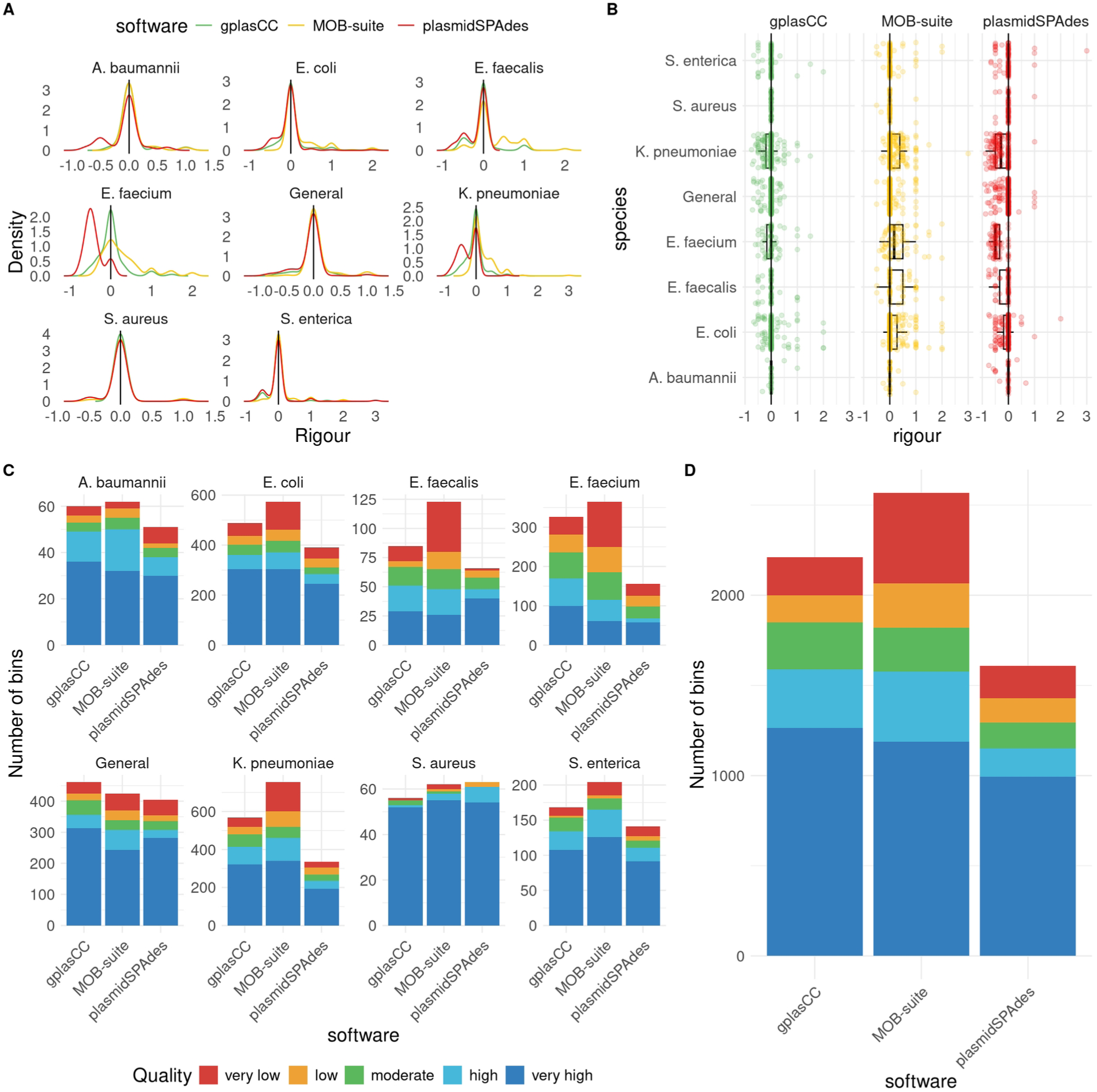
Binning rigour and binning quality. Binning rigour is defined, for each isolate, by subtracting the number of plasmids from the number of bins, divided by the number of plasmids in the isolates. Binning rigour per isolate **(A)** and binning rigour distribution per species **(B)**. Binning quality per species **(C)** and total **(D)**. Categories: Very low (completeness or accuracy less than 0.3), low (completeness or accuracy less than 0.5), moderate (completeness and accuracy less than 0.7), high (completeness and accuracy less than 0.9) and very high (completeness and accuracy higher than 0.9)

Subsequently, we separated the bins produced by each tool into quality categories based on their accuracy and completeness values (Figure 5C and D, Table 2). This showed that gplasCC had the highest percentage of bins with at least quality ‘high’ (accuracy > 0.7 and completeness > 0.7) overall. For three species and the general dataset, gplasCC had the highest absolute number of bins with ‘very high’ (accuracy > 0.90 and completeness > 0.90) quality. The exceptions were *S. enterica* and *K. pneumoniae*, where MOB-suite had a number of bins (126 and 340, respectively), *E. faecalis*, where PlasmidSPAdes had a higher number (40) and *S. aureus*, where both other tools outperformed gplasCC. Additionally, gplasCC achieved the highest percentage of bins with at least quality ‘high’ in 5 species and the general dataset (except *E. faecalis* and *S.aureus)*, and the lowest percentage of bins with quality at most ‘low’ in 5 species and the general dataset (except *A. baumannii* and *E. faecalis*).

**Table 2:** Percentage of plasmid bins in each quality category, split by tool and species.

| Software | Species | Very low | Low | Moderate | High | Very high |
| --- | --- | --- | --- | --- | --- | --- |
| MOB-suite | *A. baumannii* | 4.84% | 6.45% | 8.06% | 29.03% | 51.61% |
| gplasCC | *A. baumannii* | 6.67% | 5.00% | 6.67% | 21.67% | 60.00% |
| plasmidSPAdes | *A. baumannii* | 13.73% | 3.92% | 7.84% | 15.69% | 58.82% |
| MOB-suite | *E. coli* | 19.51% | 7.84% | 7.84% | 12.02% | 52.79% |
| gplasCC | *E. coli* | 10.66% | 6.97% | 8.40% | 11.68% | 62.30% |
| plasmidSPAdes | *E. coli* | 11.51% | 8.95% | 6.65% | 10.23% | 62.66% |
| MOB-suite | *E. faecalis* | 34.96% | 12.20% | 13.82% | 17.89% | 21.14% |
| gplasCC | *E. faecalis* | 15.29% | 5.88% | 18.82% | 25.88% | 34.12% |
| plasmidSPAdes | *E. faecalis* | 3.03% | 9.09% | 15.15% | 12.12% | 60.61% |
| MOB-suite | *E. faecium* | 31.78% | 17.53% | 19.18% | 14.79% | 16.71% |
| gplasCC | *E. faecium* | 13.80% | 13.80% | 20.55% | 21.17% | 30.67% |
| plasmidSPAdes | *E. faecium* | 19.87% | 17.31% | 19.23% | 6.41% | 37.18% |
| MOB-suite | *K. pneumoniae* | 20.29% | 10.88% | 7.82% | 15.92% | 45.09% |
| gplasCC | *K. pneumoniae* | 8.63% | 6.87% | 11.44% | 16.37% | 56.69% |
| plasmidSPAdes | *K. pneumoniae* | 8.66% | 11.34% | 10.15% | 12.24% | 57.61% |
| MOB-suite | *General* | 12.74% | 7.31% | 7.31% | 15.09% | 57.55% |
| gplasCC | *General* | 7.81% | 4.77% | 10.20% | 9.33% | 67.90% |
| plasmidSPAdes | *General* | 12.35% | 4.69% | 7.16% | 6.17% | 69.63% |
| MOB-suite | *S. aureus* | 3.23% | 1.61% | 1.61% | 4.84% | 88.71% |
| gplasCC | *S. aureus* | 1.79% | 0.00% | 3.57% | 1.79% | 92.86% |
| plasmidSPAdes | *S. aureus* | 0.00% | 3.17% | 0.00% | 11.11% | 85.71% |
| MOB-suite | *S. enterica* | 9.31% | 1.96% | 7.84% | 19.12% | 61.76% |
| gplasCC | *S. enterica* | 7.14% | 1.19% | 11.90% | 15.48% | 64.29% |
| plasmidSPAdes | *S. enterica* | 9.93% | 4.26% | 7.09% | 14.18% | 64.54% |

The ability of each tool to correctly assign plasmid-borne ARGs to predictions was evaluated (Supplementary Figure S5). gplasCC and MOB-suite performed comparably well (difference in missed ARGs below 10 percentage points) in all species except *A. baumannii, E. faecium* and *S. enterica*. gplasCC was able to correctly assign most plasmid-borne ARGs in the General dataset. In contrast, plasmidSPAdes correctly assigned lower fractions of plasmid-borne ARGs in four datasets (*E. coli*, *K. pneumoniae*, *S. enterica* and the general dataset), ranging from 43.1% (*S. enterica*) to 60.8% (*E. coli*). In *S. aureus*, all tools performed well for *S. aureus*, correctly assigning more than 85% of all plasmid-borne ARGs. However, MOB-suite and plasmidSPAdes correctly classified more than 90% of ARGs, whereas gplasCC classified a lower percentage of ARGs. However, when taking into account the number of wrongly predicted on plasmids, gplasCC includes fewer chromosomal ARGs in four datasets (*A. baumannii, E. faecium, S. aureus* and the general dataset), being second to PlasmidSPAdes in the other datasets.

We also examined whether plasmid bins were contaminated with chromosomal sequences (Supplementary Figure S6A). PlasmidSPAdes showed the largest number of contaminated plasmid bins (n=1248), with 77.9% (n=972) of these were predominantly composed of chromosomal sequences (chromosome contamination >50%). Likewise, a total of 626 plasmid predictions made by MOB-suite were contaminated, and 45.85% (n=287) of these were strictly chromosomal (chromosome contamination = 100%). gplasCC had the lowest number of contaminated bins (n=205) and most (n=108, 52.7%) showed contamination fractions below 50%. Some differences were observed when analysing each species separately (Supplementary Figure S6B). For *A. baumannii* and *E. faecium*, MOB-suite displayed the highest number of contaminated bins, while plasmidSPAdes did so in all other species.

Finally, we analysed predictions of small plasmids (n=1079). Overall, gplasCC detected 87.5% of small plasmids, surpassing MOB-suite (76.8%) and plasmidSPAdes (79.5%) (Supplementary Figure S7A). gplasCC and MOB-suite achieved accuracy medians of 1 in all species and median completeness values of 1 in all but one species (*E. faecium*). PlasmidSPAdes generally had lower accuracy and completeness values, although the lowest median accuracy and completeness values were 0.93 and 0.97, respectively (*S. aureus and S. enterica*).

### GplasCC is able to successfully assign repeated elements to the plasmidome

To assess the performance of gplasCC in correctly classifying repeats, we compared their classifications to their true origin (see methods). Metrics on repeat classification for every species are shown in Figure 6. Across all species, the count of correctly classified repeats— whether plasmidome-derived (TP) or chromosomal-origin (TN)—significantly exceeded incorrect classifications (FP and FN). Notably, in every case, the number of repeats correctly assigned to the plasmidome fraction (TP) was consistently higher than the count of chromosomal repeats misclassified into this fraction (FP).

**Figure 6.**
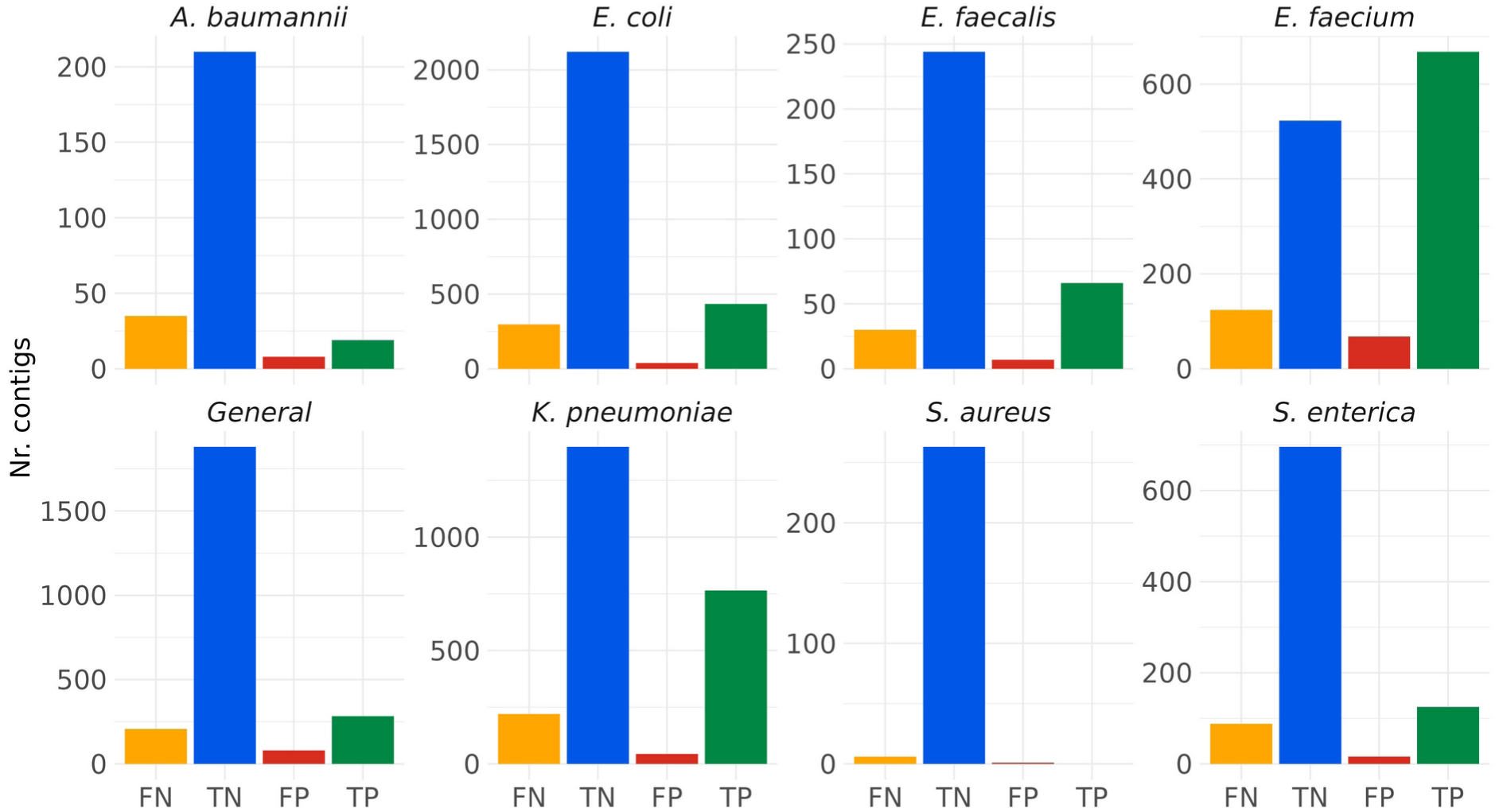
Repeat assignment of gplasCC. Repeats are considered true positive (TP) if correctly assigned to the plasmidome fraction, false positive (FP) when incorrectly assigned to this fraction, true negative (TN) when correctly not assigned to any of the plasmid bins and false negative (FN) when incorrectly not added to the plasmidome.

To further examine the accuracy of repeat assignments, we analyzed the lengths and plasmidCC scores of plasmidome repeats (Supplementary Figure S8). Correctly identified plasmidome repeats (TP) tended to be shorter, with a median length of 1,472 bp (IQR: 1,223 bp - 2,225.5 bp), compared to misclassified plasmidome repeats (FN), which had a median length of 1,787 bp (IQR: 1,323.75 bp - 4,028.5 bp). Additionally, plasmidCC scores revealed that nearly half of repeats (47.71% of TP and 50.93% of FN) were either unclassified or misclassified as chromosomal by our binary classification method, underscoring the challenges in predicting repeat origins solely based on contig DNA sequence information.

## DISCUSSION

In this work, we present gplasCC, a tool capable of reconstructing plasmids from short-read sequences across a wide array of bacterial species. Integrated within gplasCC is plasmidCC, a binary classification tool that accurately identifies plasmid contigs. We benchmarked plasmidCC against other publicly available classification tools and gplasCC against MOB-suite and plasmidSPAdes, using a dataset comprising 981 complete bacterial genomes and 2,620 plasmids from more than 100 different species. To our knowledge, this represents the largest and most diverse dataset ever utilized to benchmark plasmid reconstruction tools. Using the same dataset for both benchmarks did not compromise the validity of the study, as plasmidCC’s execution did not influence gplasCC’s performance. Furthermore, no hyperparameter tuning of either tool was performed based on the benchmarking results.

PlasmidCC outperformed other existing binary classifiers for all but one evaluated species (*S. enterica*), for which a majority voting principle of all benchmarked tools demonstrated better performance. The superior performance of plasmidCC suggests that much of the bacterial pan-plasmidome diversity is well-represented within the plasmid sequences included in the plasmidCC databases. However, this result may also reflect biases introduced by selecting sequences primarily from public databases like RefSeq and SRA, which are known to overrepresent clinical isolates and a limited range of geographical origins^45^. Incorporating sequences from understudied sources or regions could help evaluate the broader applicability of the methods described here.

A notable limitation of most classification tools is their binary output, which forces contigs to be classified as either plasmid- or chromosome-derived. This can hinder the accurate prediction of mobile genetic elements that are found on both plasmids and chromosomes^22,46,47^. To partially overcome this limitation, the Centrifuge-based classifiers described in this paper can assign contigs to an ‘unclassified’ category if they align to the plasmidome and chromosome fraction of the database in similar proportions. Despite this, ambiguous contigs often remain misclassified without examining their genomic context, as highlighted in Supplementary Figure S8. By integrating plasmidCC predictions with the repeat-resolution feature of gplasCC, we successfully addressed many of these misclassifications. Recent advancements, such as the graph neural network approach in plASgraph2^48^, enrich contigs with features from neighboring nodes, improving classification performance. We speculate that further growth in public datasets could enhance these graph-based approaches for ambiguous contig classification.

When reconstructing individual ARG-plasmids of *S. enterica* and *S. aureus*, gplasCC, MOB-suite and plasmidSPAdes performed comparably well, displaying high values of completeness and accuracy. However, gplasCC outperformed the other tools when reconstructing ARG-plasmids of *E. coli, E. faecium*, *E. faecalis*, *K. pneumoniae* and the General dataset. For these species, predictions generated by MOB-suite had lower completeness values while plasmidSPAdes predictions lacked accuracy. Furthermore, investigating the number of repeated regions in these species revealed that *E. faecium* and *K. pneumoniae* contained more repeated regions when compared to those of other species. Although repeated regions were excluded from the calculation of completeness and accuracy, these genomic characteristics are expected to cause more fragmented and entangled assemblies, which would lead to difficulties when attempting to reconstruct plasmids with tools that uniquely rely on either reference- or graph-based approaches as MOB-suite and plasmidSPAdes do. The observation that gplasCC predictions were less affected demonstrates the benefits of combining assembly graph information with accurate contig classifications to predict individual plasmids.

The balance between accuracy and completeness directly relates to a tool’s capacity to produce the correct number of plasmid bins for a genome. Low per-bin completeness suggests excessive bin fragmentation, whereas low per-plasmid accuracy indicates difficulty separating distinct plasmids into unique bins. PlasmidSPAdes’ lower accuracy highlights its tendency to group multiple plasmids into fewer bins, often contaminated with chromosomal sequences. Conversely, MOB-suite’s high accuracy reflects its precision in producing distinct bins, albeit at the cost of lower completeness. For MOB-Suite, these observations are in line with earlier findings in *E. coli* ^21,40^, showing that binning rigour is a large bottleneck in MOB-Suite’s performance. GplasCC’s integration of classification and graph-based binning achieves a favorable balance, resulting in high-quality plasmid reconstructions with minimal chromosomal contamination.

For large ARG-plasmids in underrepresented species, gplasCC demonstrated comparable accuracy to MOB-suite and higher completeness than plasmidSPAdes. Combined with its minimal chromosomal contamination, gplasCC and its species-independent Centrifuge model hold promise for exploring the pan-plasmidome diversity in publicly available datasets. Additionally, these tools may be adapted for metagenomic analyses with further computational optimizations, such as parallelization and adjustments for variable sequencing coverage. However, since the plasmidCC databases were constructed from culturable bacteria, their performance on unculturable species, often found in metagenomic samples, remains uncertain.

Despite its advancements, gplasCC has limitations. Reconstructing plasmids in isolates with highly complex plasmidomes remains challenging, particularly when multiple plasmids share repeated elements. Incorporating prior estimates of plasmid numbers, such as those derived from incompatibility groups, relaxases, or replication origins, may enhance the accuracy of plasmidome network partitioning. Additionally, refining the repeat-resolution algorithm, such as by optimizing short-walk parameters, could further improve plasmid reconstructions.

In conclusion, we demonstrated that gplasCC, in combination with a robust classifier like plasmidCC, provides the most effective method for reconstructing plasmids from short-read sequencing data across diverse bacterial species. These tools offer a powerful approach for advancing plasmidomics research and understanding AMR dissemination in bacterial populations.

## DATA AVAILABILITY

The versions of gplasCC and plasmidCC used in this study are publicly available on GitLab at https://gitlab.com/mmb-umcu/gplascc and https://gitlab.com/mmb-umcu/plasmidCC.

Additionally, both gplasCC and plasmidCC can be used as standalone tools available through the Python Package Index (pip).

Codebooks of the analyses performed in this study can be found at https://gitlab.com/jpaganini/gplascc_benchmark.

Centrifuge databases built in this study are publicly available at zenodo.org and can be downloaded from the following links:

*K. pneumoniae*: https://zenodo.org/record/7194565/files/K_pneumoniae_plasmid_db.tar.gz

*S. enterica*: https://zenodo.org/record/7133407/files/S_enterica_plasmid_db.tar.gz

*S. aureus*: https://zenodo.org/record/7133406/files/S_aureus_plasmid_db.tar.gz

*A. baumannii*: https://zenodo.org/record/7326823/files/A_baumannii_plasmid_db.tar.gz

*E. faecalis:* https://zenodo.org/records/10471306/files/E_faecalis_centrifuge_db.tar.gz

*E. faecium:* https://zenodo.org/records/10472051/files/E_faecium_centrifuge_db.tar.gz

General model: https://zenodo.org/record/7431957/files/general_plasmid_db.tar.gz

