## Supplementary Material for "gplasCC: classification and reconstruction of plasmids from short-read sequencing data for any bacterial species"

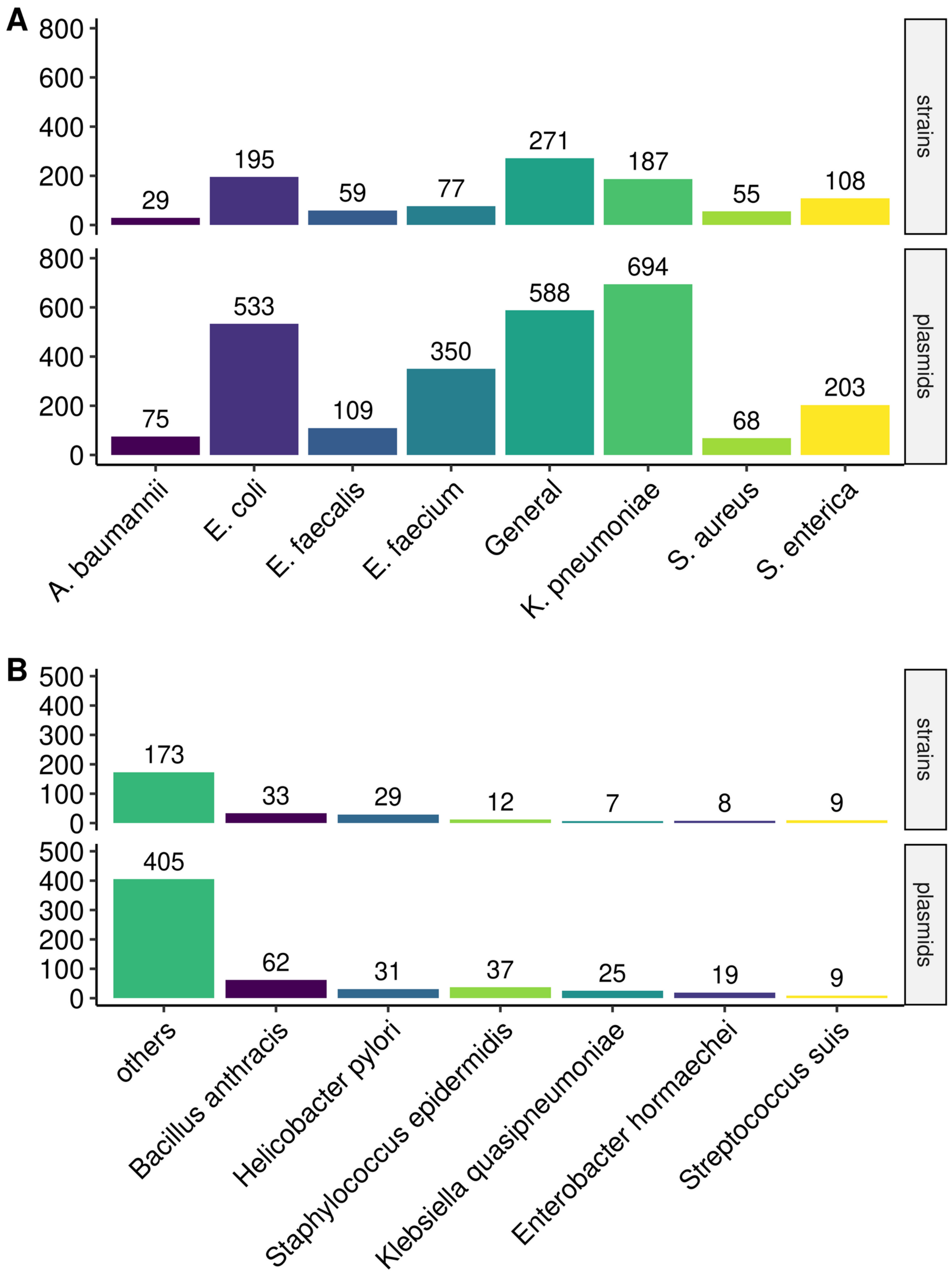


**Supplementary Figure S1.** A) Number of genomes and plasmids from common human pathogens, highly abundant in databases, included in the benchmark dataset. B) Number of genomes and plasmids of less-frequent species included in the benchmark dataset. The 'Other' category includes species with less than 10 genomes in the dataset.

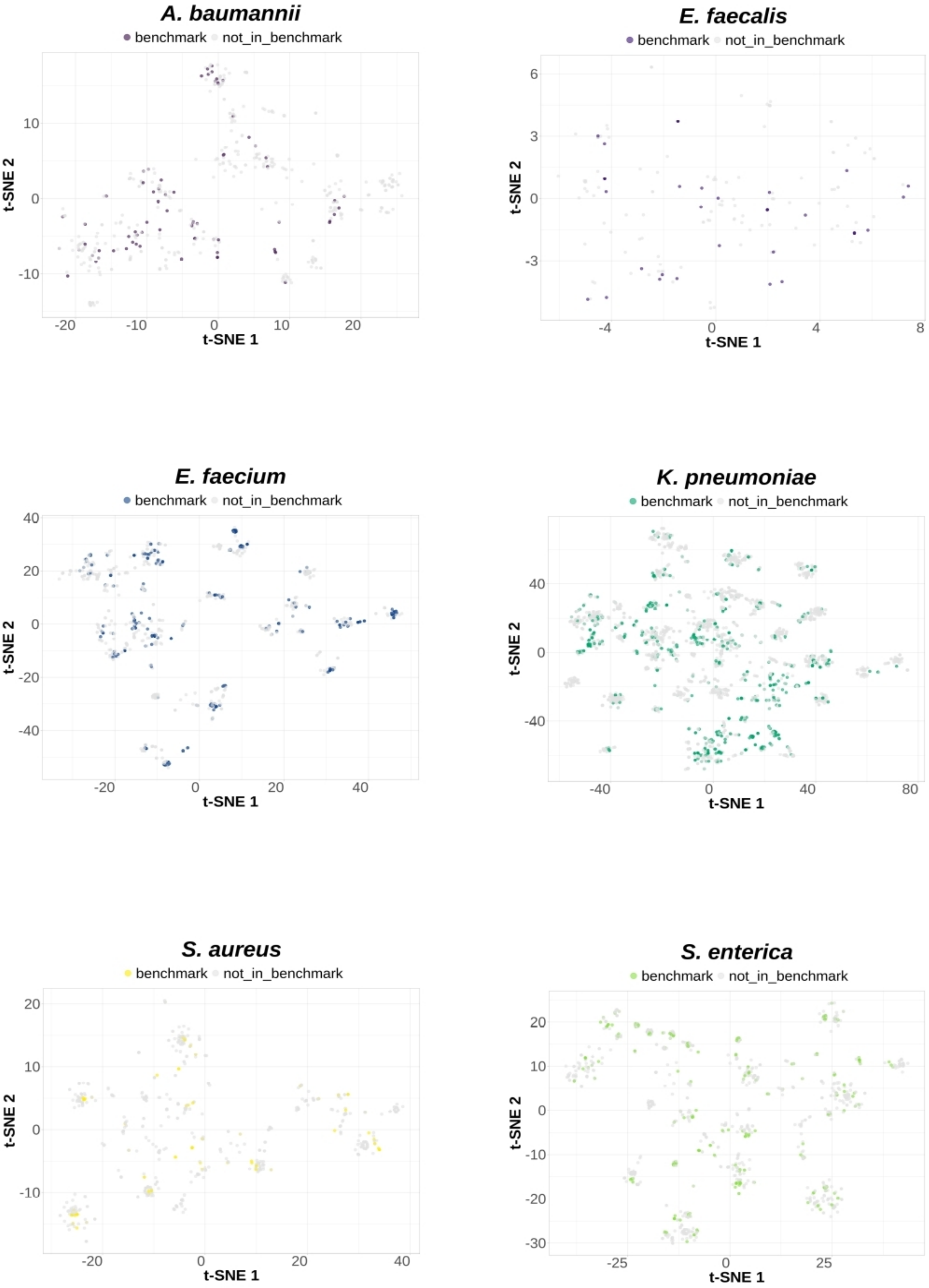


**Supplementary Figure S2.** t-SNE visualization of MASH distances (s=10.000, k=21) between all complete plasmid sequences in NCBI for *A. baumannii, E. faecalis, E. faecium, K. pneumoniae, S. aureus and S. enterica*. Plasmids that are colored are included in the benchmark datasets for the respective species-specific models.

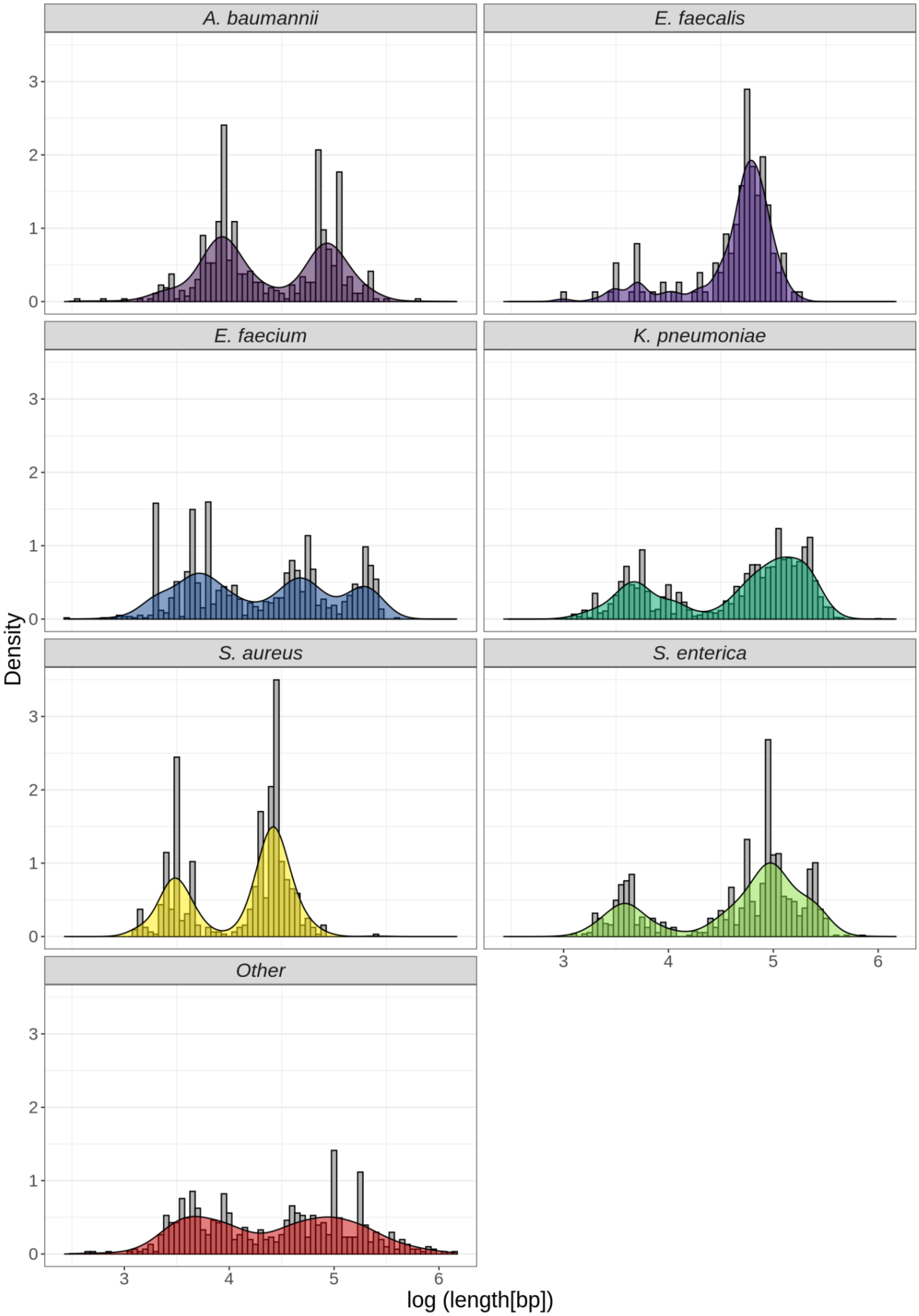


**Supplementary Figure S3.** Plasmid size distribution per species. Lengths were obtained using all complete genomes available in Public databases. The y-axis shows Kernell probability density function values, based on the abundance of the different plasmid sizes.

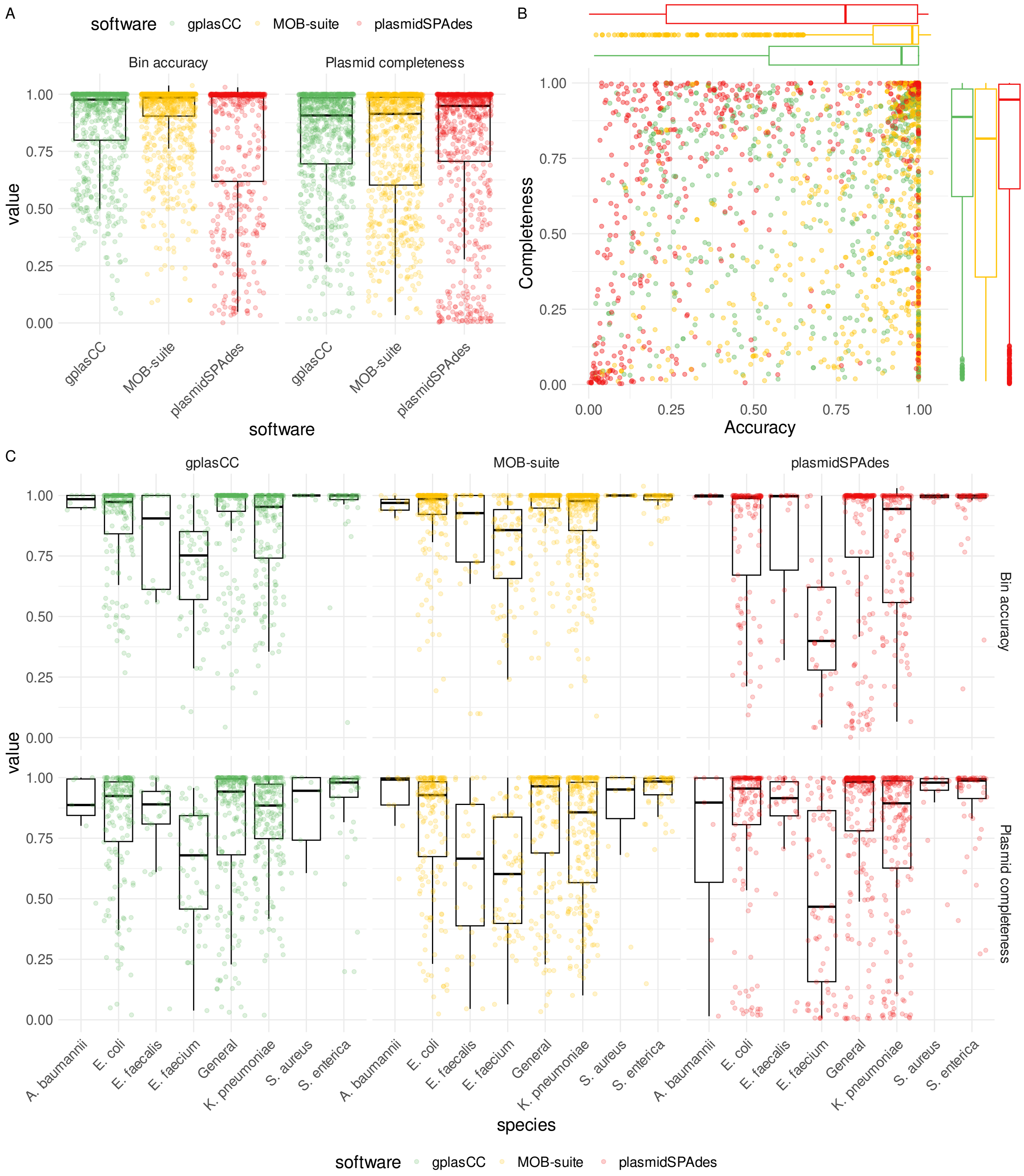


**Supplementary Figure S4.** Reconstruction metrics from gplasCC, MOB-suite and plasmidSPAdes for large plasmids that don't carry ARGs across multiple species. A) Bin accuracy, and plasmid completeness per tool, B) plasmid completeness (y-axis) vs bin accuracy (x-axis) for all plasmids. C) Plasmid completeness and bin accuracy per tool and per species. Bars indicate detected plasmids.

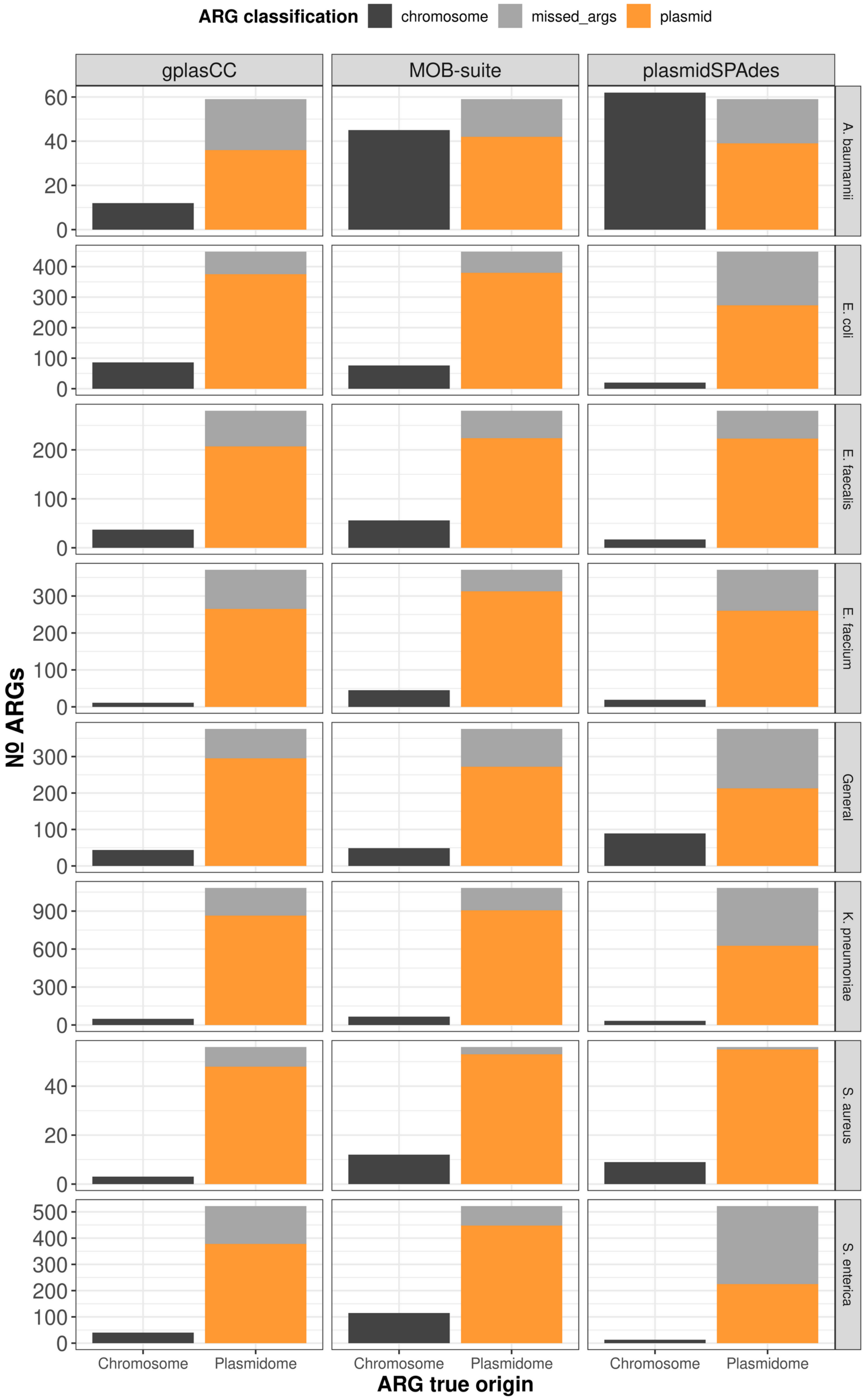


**Supplementary Figure S5.** Number of true plasmid-borne ARGs included (detected) and missing (not detected) from plasmid predictions. Chromosomal ARGs included in plasmid predictions are labelled as contamination.

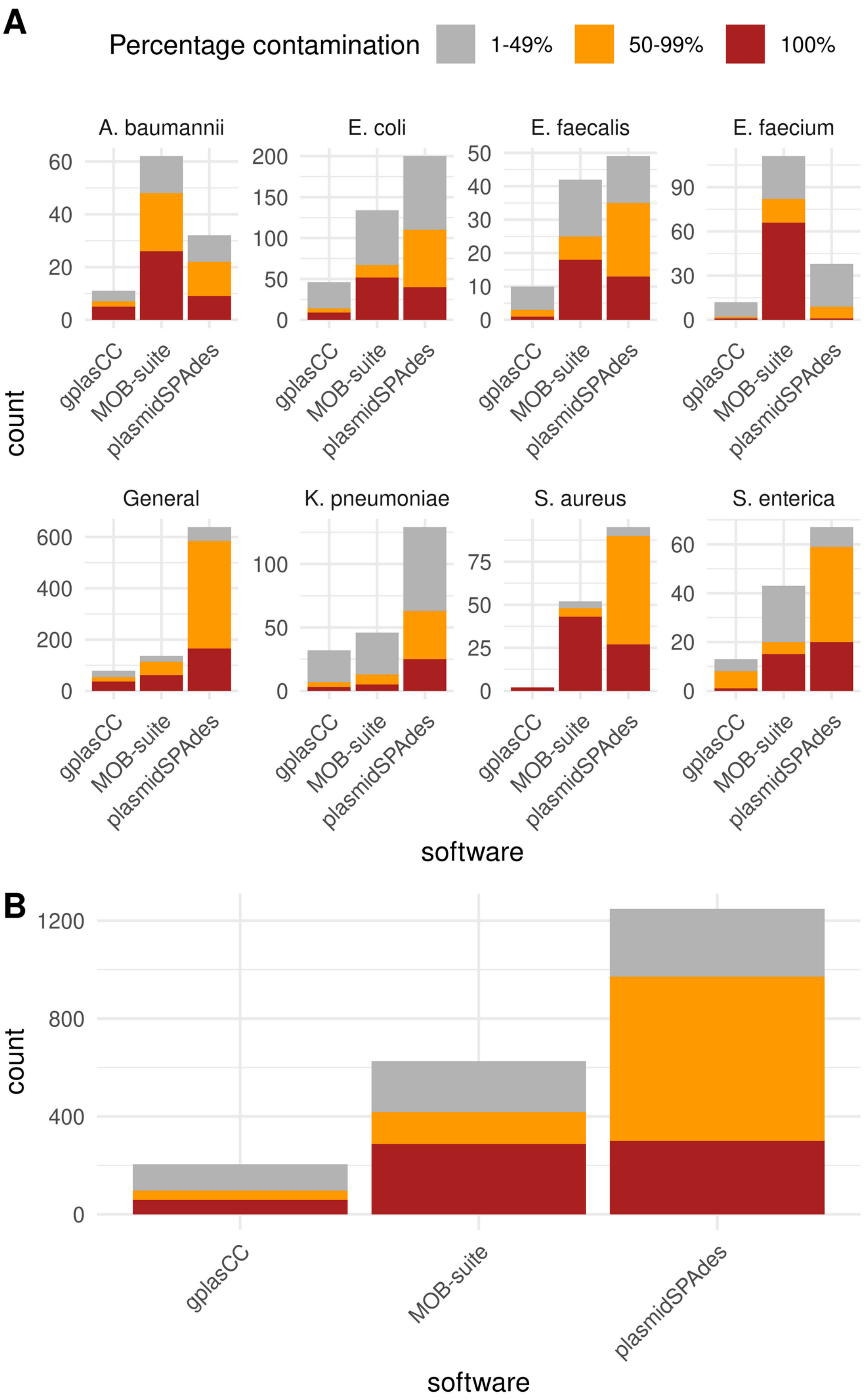


**Supplementary Figure S6.** Number of plasmid predictions that include chromosomal sequences (contamination), for all species together in (A) and individually in (B). In red, predictions that are composed solely of chromosomal sequence. In orange, predictions that contain more than 50% of chromosomal sequences in length. In grey, predictions that contain less than 50% of chromosomal sequences.

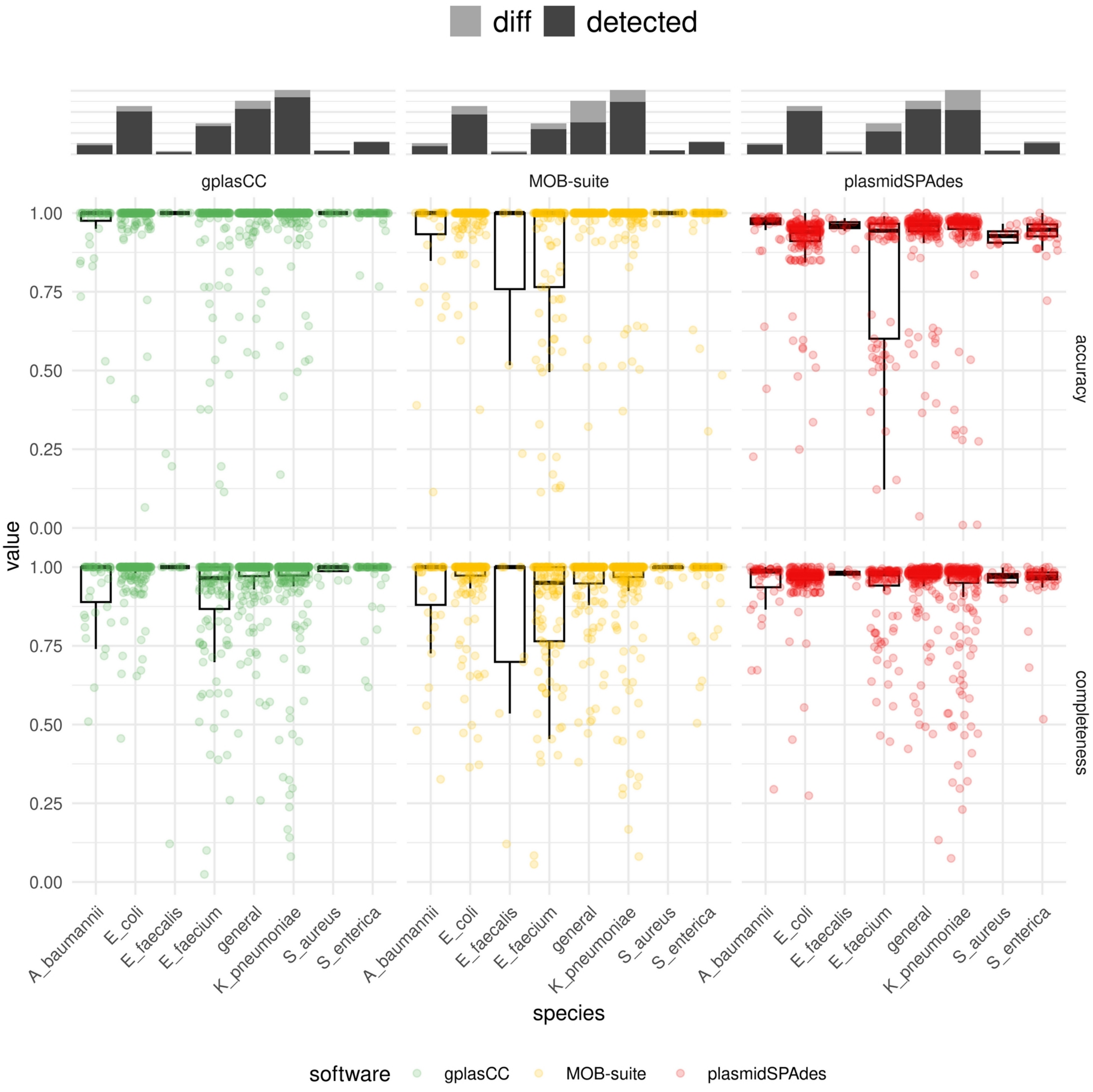


**Supplementary Figure S7**. A) Fraction of small plasmids detected (black bars) out of the total (gray bars) by each tool per species. Small plasmids are defined as those with lengths smaller than 18kb for *A.bumannii, E. coli, E. faecium, K. pneumoniae and S. enterica* and less-frequent species (general) and smaller than 8kb for *S. aureus.* B) Reconstruction metrics of small plasmids.

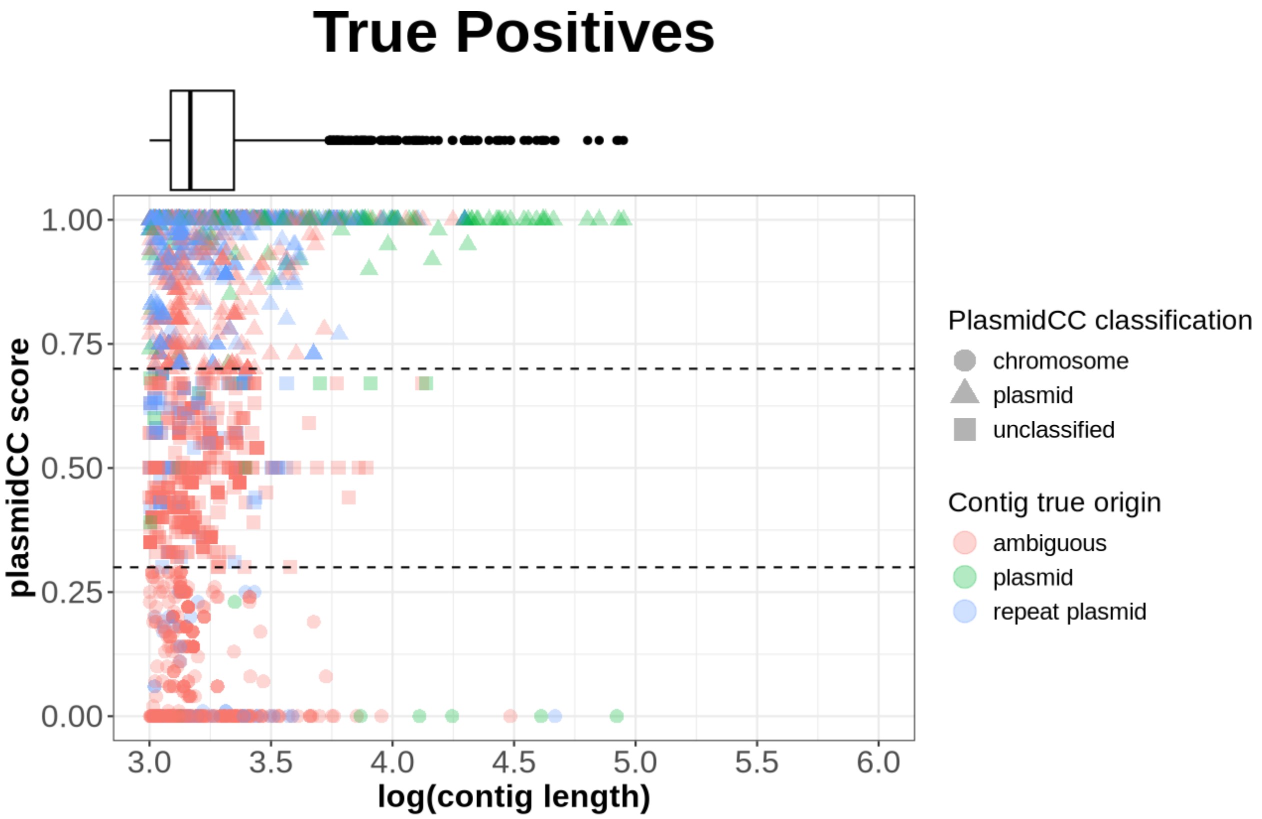


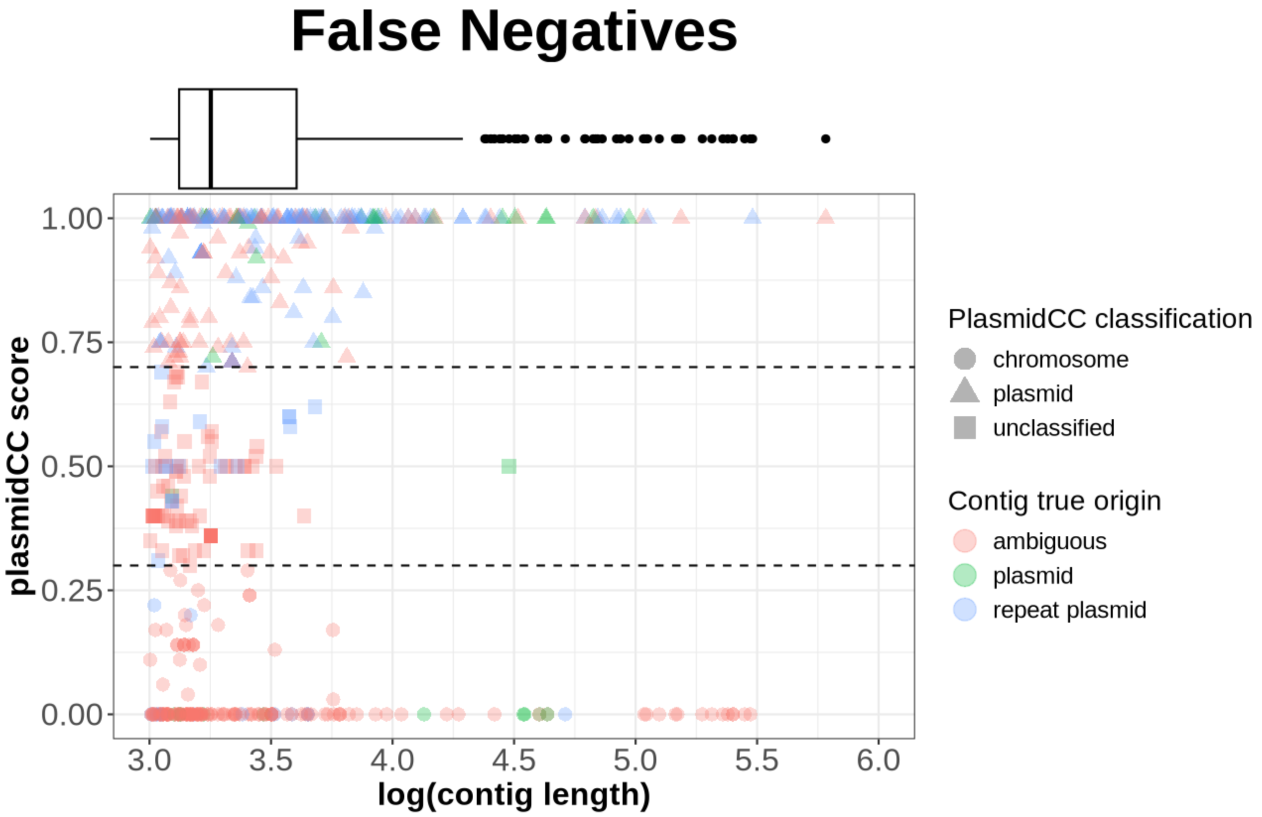


**Supplementary Figure S8.** PlasmidCC score vs contig length for ambiguous contigs correctly assigned to the plasmidome fraction (True Positives) and missed (True Negatives) by gplasCC's repeat resolution module.

**Supplementary Table S1.** Number of plasmids and ARGs per species and length classification

| Species | Length classification | Number of plasmids | Number of ARGs |
|---|---|---|---|
| *A. baumannii* | large | 22 | 56 |
| *A. baumannii* | small | 53 | 4 |
| *E. coli* | large | 305 | 444 |
| *E. coli* | small | 228 | 5 |
| *E. faecalis* | large | 94 | 277 |
| *E. faecalis* | small | 15 | 3 |
| *E. faecium* | large | 204 | 374 |
| *E. faecium* | small | 146 | 3 |
| *K. pneumoniae* | large | 390 | 1036 |
| *K. pneumoniae* | small | 304 | 14 |
| *S. aureus* | large | 49 | 53 |
| *S. aureus* | small | 19 | 4 |
| *S. enterica* | large | 142 | 498 |
| *S. enterica* | small | 61 | 25 |
| general | large | 335 | 358 |
| general | small | 253 | 20 |

**Supplementary Table S2.** 25%, 50% and 75% percentile of accuracy and completeness per species and software

| *Species* | *Software* | *Q1 bin accuracy* | *Median bin accuracy* | *Q3 bin accuracy* | *Q1 plasmid completeness* | *Median plasmid completeness* | *Q3 plasmid completeness* |
|---|---|---|---|---|---|---|---|
| *A. baumannii* | MOB-suite | 0.918 | 0.966 | 0.993 | 0.740 | 0.870 | 0.905 |
| | gplasCC | 0.929 | 0.978 | 1.000 | 0.368 | 0.791 | 0.894 |
| | plasmidSPAdes | 0.742 | 0.929 | 1.000 | 0.262 | 0.894 | 0.952 |
| *E. coli* | MOB-suite | 0.694 | 0.917 | 0.987 | 0.421 | 0.779 | 0.931 |
| | gplasCC | 0.714 | 0.933 | 0.984 | 0.575 | 0.846 | 0.933 |
| | plasmidSPAdes | 0.424 | 0.672 | 0.997 | 0.365 | 0.793 | 0.949 |
| *E. faecalis* | MOB-suite | 0.877 | 1.000 | 1.000 | 0.487 | 0.786 | 0.958 |
| | gplasCC | 0.851 | 0.959 | 1.000 | 0.715 | 0.857 | 0.937 |
| | plasmidSPAdes | 0.638 | 0.960 | 1.000 | 0.849 | 0.944 | 0.968 |
| *E. faecium* | MOB-suite | 0.656 | 0.839 | 0.999 | 0.378 | 0.600 | 0.728 |
| | gplasCC | 0.746 | 0.858 | 0.927 | 0.586 | 0.694 | 0.753 |
| | plasmidSPAdes | 0.266 | 0.385 | 0.592 | 0.150 | 0.729 | 0.817 |
| *K. pneumoniae* | MOB-suite | 0.894 | 0.963 | 1.000 | 0.589 | 0.814 | 0.900 |
| | gplasCC | 0.697 | 0.934 | 0.974 | 0.650 | 0.831 | 0.918 |
| | plasmidSPAdes | 0.583 | 0.909 | 0.992 | 0.405 | 0.849 | 0.920 |
| *S. aureus* | MOB-suite | 0.950 | 1.000 | 1.000 | 0.989 | 1.000 | 1.000 |
| | gplasCC | 1.000 | 1.000 | 1.000 | 0.986 | 1.000 | 1.000 |
| | plasmidSPAdes | 0.986 | 0.988 | 0.991 | 0.992 | 0.994 | 0.995 |
| *S. enterica* | MOB-suite | 0.899 | 0.967 | 1.000 | 0.776 | 0.919 | 0.958 |
| | gplasCC | 0.793 | 0.961 | 1.000 | 0.795 | 0.908 | 0.956 |
| | plasmidSPAdes | 0.814 | 0.970 | 1.000 | 0.414 | 0.861 | 0.944 |
| general | MOB-suite | 0.866 | 0.968 | 1.000 | 0.586 | 0.806 | 0.917 |
| | gplasCC | 0.779 | 0.955 | 1.000 | 0.728 | 0.871 | 0.950 |
| | plasmidSPAdes | 0.605 | 0.902 | 0.996 | 0.766 | 0.890 | 0.950 |
